# Gait speed and individual characteristics can be used to predict specific gait metric magnitudes in neurotypical adults

**DOI:** 10.1101/2022.10.27.513226

**Authors:** Maryana Bonilla Yanez, Sarah A. Kettlety, James M. Finley, Nicolas Schweighofer, Kristan A. Leech

## Abstract

**Background:** Gait biofeedback is commonly used to reduce gait dysfunction in a variety of clinical conditions. In these studies, participants alter their walking to reach the desired magnitude of a specific gait parameter (the biofeedback target) with each step. Biofeedback of parameters such as anterior ground reaction force and step length have been well-studied. Yet, there is no standardized methodology to set the target magnitude of these parameters. Here we present an approach to predict the anterior ground reaction force and step length of neurotypical adults walking at different speeds as a potential method for personalized gait biofeedback.

**Research question:** To determine if anterior ground reaction force and step lengths achieved during neurotypical walking could be predicted using gait speed and participants’ demographic and anthropomorphic characteristics.

**Methods:** We analyzed kinetic and kinematic data from 51 neurotypical adults who walked on a treadmill at up to eight speeds. We calculated the average peak anterior ground reaction force and step length of the right lower extremity at each speed. We used linear mixed-effects models to evaluate the effect of speed, leg length, mass, and age on anterior ground reaction force and step length. We fit the model to data from 37 participants and validated predictions from the final models on an independent dataset from 23 participants.

**Results:** Final prediction models for anterior ground reaction force and step length both included speed, speed squared, age, mass, and leg length. The models both showed strong agreement between predicted and actual values on an independent dataset.

**Significance:** Anterior ground reaction force and step length for neurotypical adults can be predicted given an individual’s gait speed, age, leg length, and mass. This may provide a standardized method to personalize targets for individuals with gait dysfunction in future studies of gait biofeedback.

## Introduction

Gait training is a common component of rehabilitation paradigms for individuals with gait dysfunction secondary to a variety of musculoskeletal and neurological conditions. This training typically involves interventions that target stability, endurance, speed, and movement patterns during walking [1–5]. In this context, clinicians often provide external cues or feedback to change different aspects of walking that are associated with gait speed [6,7], fall risk [8–10], orthopedic injury [11,12], and metabolic cost [6,10,13,14] in many patient populations. This has prompted research studies designed to evaluate the use of real-time biofeedback to change gait biomechanics.

These studies have largely focused on the effects of visual gait biofeedback, with kinematic and kinetic gait parameters most often fed back to the learners [15]. In studies of gait dysfunction post-stroke, for example, visual biofeedback is often used to promote increases in step length (and reductions in step length asymmetry) [16–19] or increases in peak anterior ground reaction force [20,21]. These gait biofeedback parameters are also common targets in studies of gait in older adults [22,23] and individuals with Parkinson’s Disease [24,25].

However, recent literature reviews have highlighted that the results across studies of gait biofeedback are inconsistent (for review see van Gelder, et al. 2018 [15] and Spencer, et al. 2021 [26]); due, in part, to methodological heterogeneity. They found that studies differ in the modality of the biofeedback provided and the gait parameter fed back to the learner. Even between studies that are matched for these factors, the methods to set the target values for biofeedback schemes vary. Step length biofeedback targets for people post-stroke have been set in a variety of ways. Some studies ask participants to lengthen their shorter step to the length of their longer step [18], while others instruct participants to make the right and left step lengths equal without any direction on which step length they should change [27,28]. Anterior ground reaction force biofeedback targets are often set as a percent increase from baseline walking values, but the magnitude of this increase varies between studies [22,29]. In addition to these differences, using baseline gait behavior to anchor biofeedback targets introduces the potential for ceiling effects with gait biofeedback interventions.

Here, we developed prediction models for the peak anterior ground reaction force and step length of neurotypical adults when considering walking speed, leg length, mass, and age and validated these models on an independent dataset. Models that predict the magnitude of a neurotypical peak anterior ground reaction force and step length can be leveraged as a standard methodology to set individualized biofeedback targets in gait rehabilitation research or clinical gait training.

## Methods

We completed a secondary analysis of two previously collected datasets. One to create the prediction models (a training dataset) and the other to validate them (an independent validation dataset).

### Training dataset

To train the models, we used a publicly available dataset of 42 neurotypical adults walking on a treadmill at eight different speeds, ranging from 40-145% of their self-selected speed (0.36 – 2.23 m/s) [30]. At each speed, participants walked for 90 seconds, and data from the last 30 seconds of each trial were used in this analysis, to allow the participant to adjust to each speed before recording their data. Kinematic data were recorded at 150 Hz and kinetic data at 300 Hz. To ensure accurate kinetic data, the data from four participants in the original dataset who used treadmill handrails while walking were removed. Another participant from this dataset was removed due to an aberrant offset in the force data. Therefore, we trained the models with data from a total of 37 neurotypical adults (age: 40 ± 17 years (21-71 years); leg length: 0.78 ± 0.07 meters (0.65-0.91 meters); mass: 67 ± 12 kg (45-95 kg)). For ten of the 37 participants, data from only six or seven (out of eight) trials were included in the analysis. Six of these participants were unable to complete the fastest walking speed. In the other four cases, the participants completed all walking trials, but we excluded the data from the trials in which the participants drifted across the midline of the instrumented treadmill (i.e., stepped on both the right and left force plates simultaneously). We were able to use the remaining data from these ten participants in our analysis because the statistical approach we employed (described below) is robust to missing data.

### Independent validation dataset

The independent dataset used to validate the final models consisted of 23 neurotypical adults (age: 61 ± 15 years (24-77 years); leg length: 0.78 ± 0.06 meters (0.67-0.88 meters); mass: 72 ± 18 kg (37-103 kg)) walking at their self-selected speed (0.48 to 1.29 m/s) for 2-5 minutes. These data were collected as part of another study, and the kinetic and kinematic data collection procedures have been previously reported [31]. Kinematic data were recorded at 100 Hz and kinetic at 1000 Hz. To keep the amount of data used for analysis consistent between the datasets, we extracted the first 90 seconds of each participant’s walking trial and used the last 30 seconds of that bin for analysis.

To account for the influence of other individual characteristics on peak anterior ground reaction force and step length, we extracted each participant’s age, leg length, and mass. We defined leg length as the vertical distance from the greater trochanter marker to the lateral malleolus marker while the participant stood upright. Please see the previous publications [30,31] for additional information about the data collection procedures.

### Data processing

For both datasets, we analyzed the kinematic and kinetic data from the right lower extremity using MATLAB R2021a. For the training dataset [30], kinematic and kinetic data were lowpass filtered with a cutoff frequency of 6 Hz. For the validation dataset [31], kinematic data were lowpass filtered with a cutoff frequency of 6 Hz, while kinetic, at 20 Hz [32]. Vertical ground reaction forces were then used to identify kinetic gait events. Heel strike was identified at the point when vertical ground reaction force reached 100 N and toe-off at less than 100 N.

We defined peak anterior ground reaction force as the maximum anterior ground reaction force between heel strike and toe-off. The average peak anterior ground reaction force over each 30-second trial was used for all analyses. Step length was defined as the fore-aft difference between the right and left lateral malleoli markers at heel strike. The average step length over each 30-second trial was used for all analyses.

### Statistical analyses

We used linear mixed-effects models to determine if peak anterior ground reaction force and step length magnitudes were associated with speed (m/s), mass (kg), leg length (m), and age (years). Given the known relationship between gait speed and both peak anterior ground force and step length [33], the minimum model included a fixed effect for speed. Because visual inspection of the data indicated a non-linear relationship between speed and peak anterior ground reaction force and step length, we also considered models with a quadratic transformation of speed. In addition to these fixed effects, we included both a random intercept and a random slope for speed to account for repeated measures and between-subjects variability. The best subset model selection approach was used to identify the best model in the training data set. Best subset selection was performed by first comparing all possible models with the same number of variables and selecting the one with the highest R^2^ value to yield a candidate set of six models [34]. Then models with different numbers of variables were compared using the Akaike information criterion (AIC) and the model with the lowest AIC was selected as the final prediction model. This process was completed independently for peak anterior ground reaction force and step length.

We checked for linearity, multicollinearity, outliers, and influential points for both peak anterior ground reaction force and step length prediction models. The residuals vs fitted plots showed that, for the best models, there was no deviation from linearity. The variance inflation factors (VIF) for each of the coefficients were less than 5, indicating no major issues with collinearity. We identified two significantly influential points (one walking speed for two participants) in the peak anterior ground reaction force model using Cook’s distance at a threshold of 0.12. We removed these two influential points (0.7% of the total number of data points) and refit the final peak anterior ground reaction force model. Unlike for inference, the normality and homoscedasticity assumptions of linear mixed-effects modeling do not need to be met for prediction. For the same reason, we did not select the predictors based on p-values, as the best subset model selection process determines the model that provides the best prediction versus determining which predictors are most important [34].

Next, we used an independent dataset (N = 23) to test the predictive ability of the peak anterior ground reaction force and step length models. We calculated predicted values for peak anterior ground reaction force and step length using the final models with only fixed-effects terms as the mean of the random term estimates from mixed-effects linear models is approximately zero [35]. We then calculated the R^2^ values and the root-mean-square error (RMSE) for the predicted versus actual peak anterior ground reaction force and step length. Because we found that some of the participants in the independent dataset walked with step lengths shorter than those represented in the training data, we also evaluated the R^2^ values and the RMSE of the predicted step lengths on a subset of the participants (n=18) who’s actual step lengths were represented in the training dataset.

## Results

### Model selection

The best models for each number of variables included and their AIC scores are shown in Table 1 for peak anterior ground reaction force and Table 2 for step length.

**Table 1.** Best Subset Selection for Peak Anterior Ground Reaction Force Model. The last step of the best subset selection process is shown. Comparisons across models of an increasing number of variables. The final model chosen had the lowest AIC, highlighted with (*). AGRF, anterior ground reaction force; ID, Participant Identification; SPEED_2, speed-squared;

| Model | AIC |
| --- | --- |
| AGRF ~ SPEED + (1 \| ID) | 2419.3 |
| AGRF ~ SPEED + (SPEED \| ID) | 2217.6 |
| AGRF ~ SPEED + SPEED_2 + (SPEED \| ID) | 2120.2 |
| AGRF ~ SPEED + MASS + SPEED_2 + (SPEED \| ID) | 2119.8 |
| AGRF ~ SPEED + AGE + MASS + SPEED_2 + (SPEED \| ID) | 2120.4 |
| AGRF ~ SPEED + AGE + LEGLength + MASS + SPEED_2 + (SPEED \| ID)* | 2111.7 |

**Table 2.** Best Subset Selection for Step Length Model. The last step of the best subset selection process is shown. Comparisons across models of an increasing number of variables. The final model chosen had the lowest AIC, highlighted with (*). SL, step length; ID, Participant Identification; SPEED_2, speed-squared;

| Model | AIC |
| --- | --- |
| SL ~ SPEED + (1 \| ID) | -1105.4 |
| SL ~ SPEED + (SPEED \| ID) | -1163.9 |
| SL ~ SPEED + SPEED_2 + (SPEED \| ID) | -1324.5 |
| SL ~ SPEED + LEGLENGTH + SPEED_2 + (SPEED \| ID) | -1326.0 |
| SL ~ SPEED + LEGLENGTH + MASS + SPEED_2 + (SPEED \| ID) | -1325.5 |
| SL ~ SPEED + AGE + LEGLENGTH + MASS + SPEED_2 + (SPEED \| ID)* | -1331.1 |

### Prediction of Peak Anterior Ground Reaction Force Magnitude

The model that best predicted peak anterior ground reaction force (AGRF) magnitude included fixed effects for speed, speed^2^, age, mass, and leg length, as well as a random slope (speed) and intercept. Figure 1A displays examples of individual model fits for the training data. Figure 1B shows the mean fixed-effects model (Equation 1) displayed against individual training data.

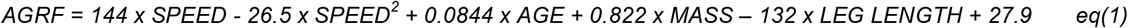

**Figure 1.**
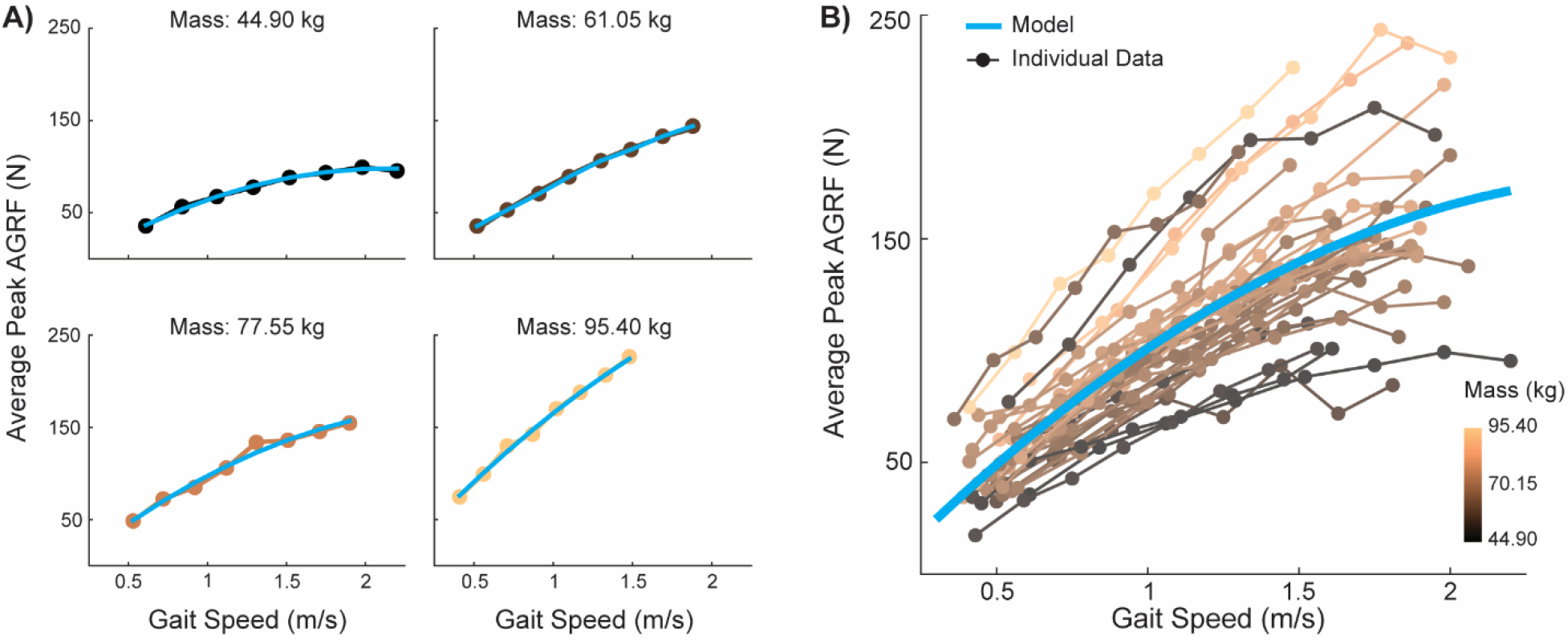
Model of peak anterior ground reaction force displayed against individual data. **A)** Representative individual model fits with fixed and random effects of data from participants of different masses. Since we included a random slope for speed and a random intercept, the individual model fits varied in steepness and intercept, depending on the participant. **B)** Mean fixed effects model of peak anterior ground reaction force (blue line) plotted with all individual data. This model was visualized through simulated fixed effect values based on the original data. Specifically, we created a vector of 20 speeds ranging from the minimum to the maximum speeds and found the age, mass, and leg length group means. These simulated values were used with the fixed-effects only model to calculate predicted values that are plotted in blue. Individual data are plotted on a color spectrum from light orange to dark brown to represent participant mass as a third dimension of the data.

agreement with the actual values of peak anterior ground reaction force magnitude in an independent dataset (N = 23; Figure 2; R^2^ relative to the unity line = 0.76). These predictions had an RMSE = 20.7 N.

**Figure 2.**
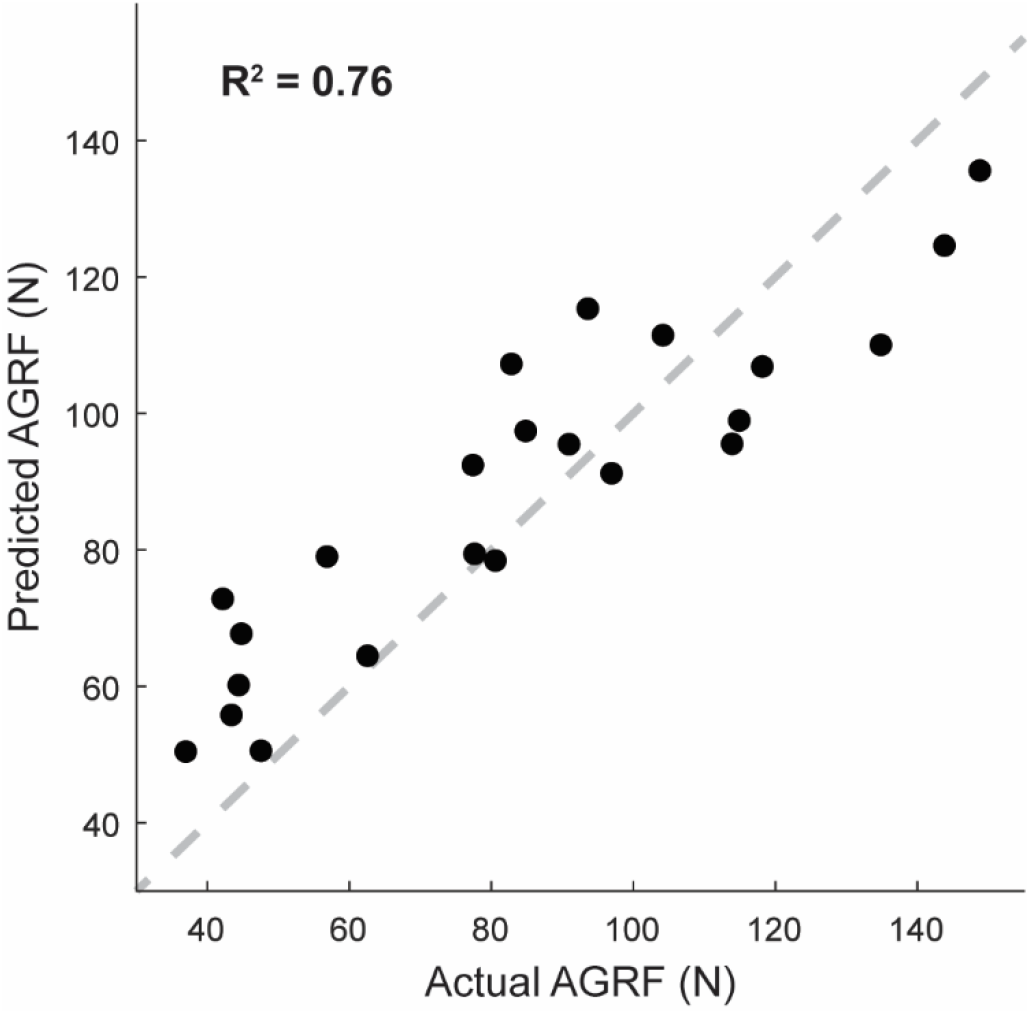
Testing the peak anterior ground reaction force (AGRF) prediction model on an independent data set. Values predicted by the model were in strong agreement with the actual values of peak anterior ground reaction force from the independent data set. The unity line is displayed as the gray dashed line.

### Prediction of Step Length Magnitude

The model that included fixed effects for speed, speed^2^, age, mass, and leg length as well as a random slope (speed) and intercept best-predicted step length (SL) magnitude. Example individual model fits on training data are displayed in Figure 3A and the mean fixed-effects model (Equation 2) against all individual training data are in Figure 3B.

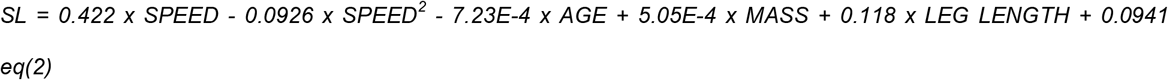

**Figure 3.**
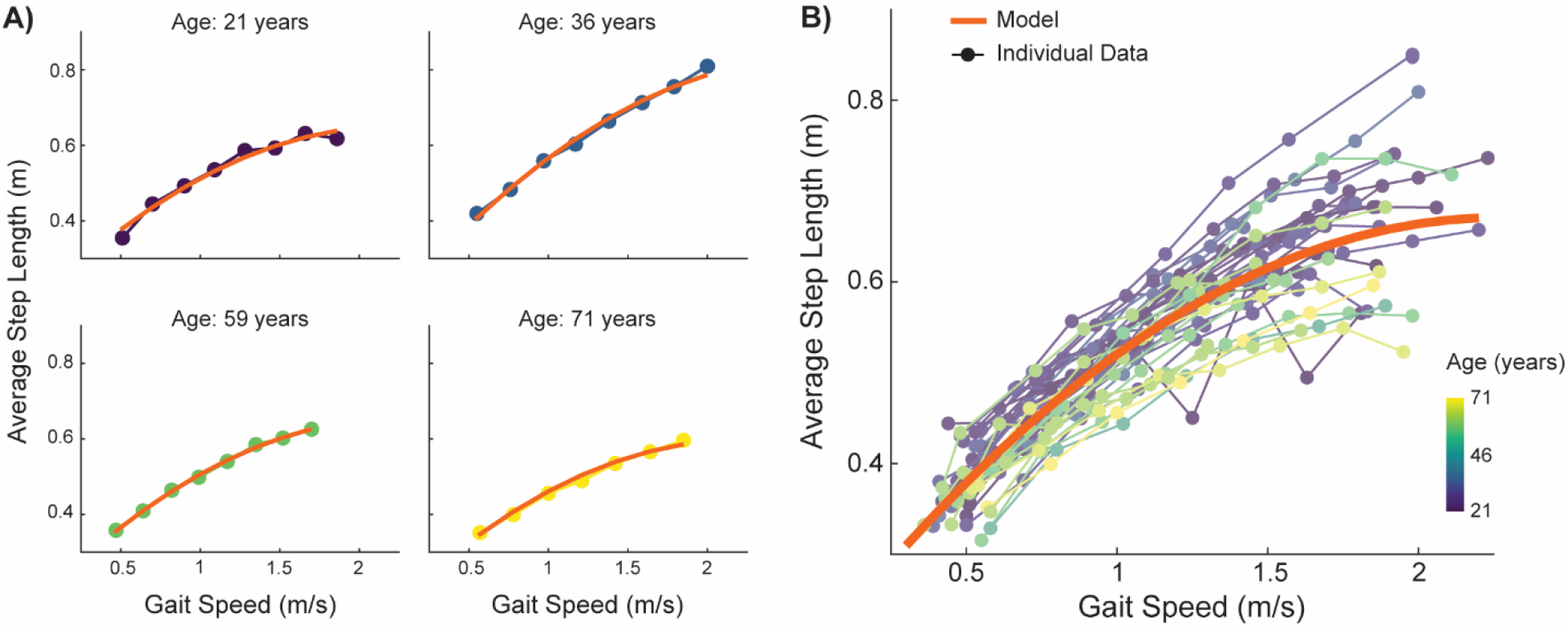
Step length model displayed against individual step length data. **A)** Representative individual model fits with fixed and random effects across the age spectrum. Since we included a random effects term for speed and a random intercept, the individual models varied in steepness and intercept, depending on the participant. **B)** Mean fixed effects model of step length (orange line) plotted with all individual data. This model was visualized with the same methods described in Figure 1B. Individual data are plotted in colors ranging from yellow to green to blue to represent each participant’s age.

Predictions from this model of step length showed moderate agreement with the actual values of step length magnitude within the full independent dataset (N = 23; Figure 4a; (R^2^ relative to the unity line = 0.69). These predictions had an RMSE = 0.09 m. Of note, the model overestimated the actual step lengths that were <0.31m, which were not represented in the training dataset.

**Figure 4.**
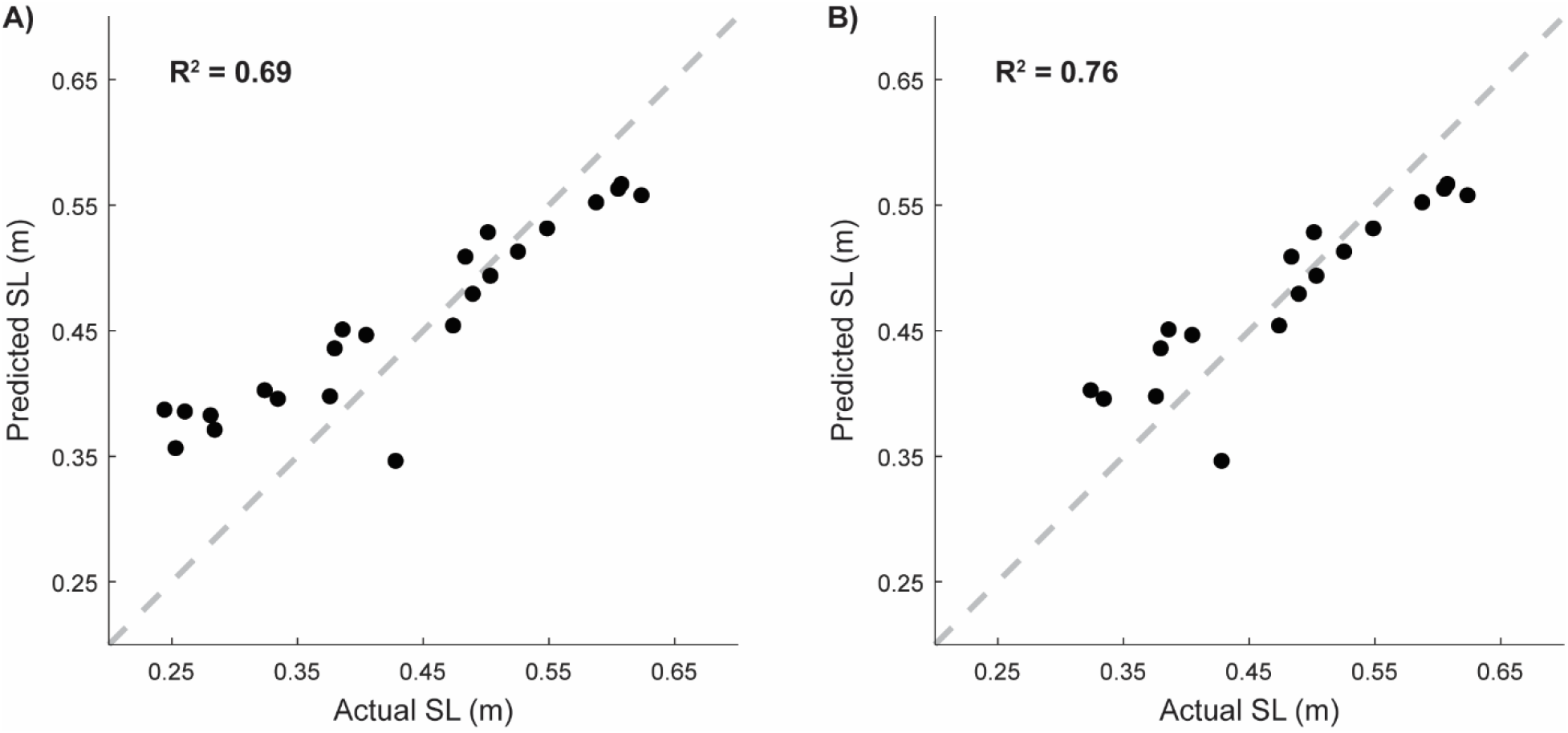
Testing the step length (SL) prediction model on an independent data set. **A)** Predicted values from the model were in moderate agreement with the actual step length magnitudes of the entire independent data set (N=23). **B)** The agreement between the predicted and actual step length values improved when we predicted only the step length values that were within the range of the step lengths represented in the training data set (N=18). The gray dashed line in both plots represents the unity line.

Because four of the participants within the independent validation set exhibited step lengths that were below the range of step lengths in the training set, we performed the same analysis on a subset of the participants who exhibited step lengths within the range of step lengths in the training set (0.31-0.85m; N=18). Step length predictions for this subset of participants showed strong agreement with the actual values (N = 18; Figure 4b; R^2^ relative to the unity line = 0.76) and had a smaller RMSE of 0.05 m.

Spreadsheets to calculate the predicted magnitudes of peak anterior ground reaction force and step length for an individual using the above equations are provided in the Supplementary Material.

## Discussion

In this study, we developed models to predict the magnitudes of peak anterior ground reaction force and step length observed in neurotypical individuals while walking at different speeds. Both models accounted for the individual’s gait speed, age, mass, and leg length. We then tested the performance of these prediction models on an independent dataset. The model-predicted magnitudes of peak anterior ground reaction force and step length were in strong and moderate agreement with the actual values of these measures, respectively. Agreement between the predicted and actual values of step length improved (from R^2^ = 0.69 to 0.76) when the step length model was used to predict step length values that were represented in the training data set. This suggests that these prediction models can generate reasonable estimates of individualized reference values for peak anterior ground reaction force and step length during walking. This has important implications for gait rehabilitation research or clinical gait training – the model equations can be used to calculate peak anterior ground reaction force and step length magnitudes that reflect a premorbid, non-pathological walking pattern given someone’s leg length, mass, age, and gait speed.

One important implication of this work is that predicted peak anterior ground reaction force and step length values can be evaluated relative to the actual values in a population with gait dysfunction. These can then be used to determine the presence and magnitude of gait pattern deficits – without the need for neurotypical control data that are demographically matched. This may significantly reduce the data collection burden for research studies and help inform clinical decision-making. These findings build upon previous studies that have developed methods to predict lower extremity joint angles and moments at different gait speeds [36–39]. In addition to predicting more clinically relevant gait metrics, our work also highlights the value of including demographic characteristics in the prediction models to attain strong agreement with an independent dataset.

The ability to generate individualized reference values for peak anterior ground reaction force and step length provides a promising standard methodology to inform individualized gait biofeedback targets. To date, biofeedback targets for anterior ground reaction force and step length have largely been anchored to baseline walking patterns [18,22]. This is largely because targets anchored to baseline behavior are individualized and assumed to be achievable. However, it is possible that this approach is masking additional capacity for change in movement patterns, particularly in cases where individuals have low baseline values for step length and anterior ground reaction force. We posit that using the values predicted from neurotypical gait behavior to inform biofeedback targets would prevent the possibility of a ceiling effect in studies of gait biofeedback. Though future work is necessary to determine the most appropriate way to implement the use of biofeedback targets based on neurotypical gait behavior to ensure the movement goals are still attainable.

There are a few limitations to this study. First, the models are trained to predict a specific range of peak anterior ground reaction force and step length values and predictions outside of this range will be less accurate – as demonstrated by the larger RMSE of the step length predictions from the full independent validation dataset compared to that from the subset of data within the training set range. In addition to this, we intentionally focused on predicting the gait metrics that are commonly studied and targeted with biofeedback in neurologic patient populations. However, other metrics may be of more interest or relevance in other diagnoses or gait pathologies (e.g., stance time asymmetry in individuals with lower extremity amputations [42]).

In conclusion, we developed prediction models for peak anterior ground reaction force and step length that demonstrated strong agreement when validated on an independent dataset. The best prediction models included terms that captured gait speed and demographic characteristics (i.e., age, mass, leg length). These models (and the associated calculation tables provided in the supplementary material) can be used to create individualized reference values of peak anterior ground reaction force and step length for use in gait rehabilitation research and clinical practice.

## Acknowledgments

This work was supported by the Magistro Family Foundation Research Grant from the Foundation for Physical Therapy Research (KAL), the National Institute of Child and Human Development (R03 HD104217-01 to KAL, 3R03HD104217-01S1 to MBY and KAL, and R01 HD091184 to JMF), the National Institute of Aging (K01 AG073467-01 to KAL), and the National Institute of Neurologic Disorders and Stroke (R21 NS120274 to NS).

